# Adaptation to life on land at 21% O_2_ via transition from ferredoxin- to NADH-dependent redox balance

**DOI:** 10.1101/680934

**Authors:** SB Gould, SG Garg, M Handrich, S Nelson-Sathi, N Gruenheit, AGM Tielens, WF Martin

## Abstract

Pyruvate:ferredoxin oxidoreductase (PFO) and iron only hydrogenase ([Fe]-HYD) are common enzymes among eukaryotic microbes that inhabit anaerobic niches. Their function is to maintain redox balance by donating electrons from food oxidation via ferredoxin (Fd) to protons, generating H_2_ as a waste product. Operating in series, they constitute a soluble electron transport chain of one-electron transfers between FeS clusters. They fulfill the same function — redox balance — served by two electron-transfers in the NADH- and O_2_-dependent respiratory chains of mitochondria. Although they possess O_2_-sensitive FeS clusters, PFO, Fd and [Fe]-HYD are also present among numerous algae that produce O_2_. The evolutionary persistence of these enzymes among eukaryotic aerobes is traditionally explained as enabling facultative anaerobic growth. Here we show that algae express enzymes of anaerobic energy metabolism at ambient O_2_ levels (21% v/v), *Chlamydomonas reinhardtii* expresses them with diurnal regulation. High O_2_ environments arose on Earth only some ∼450 million years ago. Gene presence absence and gene expression data indicate that during the transition to high O_2_ environments and terrestrialization, diverse algal lineages retained enzymes of Fd-dependent one-electron based redox balance, while the land plant and land animal lineages underwent irreversible specialization to redox balance involving the O_2_-insensitive two-electron carrier NADH.

**Highlights:**

- Algae express enzymes of anaerobic metabolism in 21% [v/v] O_2_ atmosphere, independent of anaerobiosis
- Retention of a plastid-encoded NADH dehydrogenase-like (NDH) was likely a prerequisite for the transition to life on land
- Terrestrialization and adaption to high O_2_ is accompanied by a shift to redox balance at higher midpoint potentials
- Eukaryotes adapted to high O_2_ life on land via specialization to two-electron based redox balance

## Introduction

Molecular oxygen (O_2_) had far reaching impact on evolution. From about 2.7–2.5 billion years ago onwards, cyanobacteria started using H_2_O as the electron donor for a photosynthetic electron transport chain consisting of two photosystems connected in series (Allen 2005; Fischer et al. 2016), generating O_2_ as a waste product of primary production. Before that, all life was anaerobic (Bekker et al. 2004, Rasmussen et al. 2013). Yet oxygenation of the planet did not occur quickly, as atmospheric oxygen concentrations remained low for almost 2 billion years (Lyons et al. 2014) (Fig. 1).

**Fig. 1.**
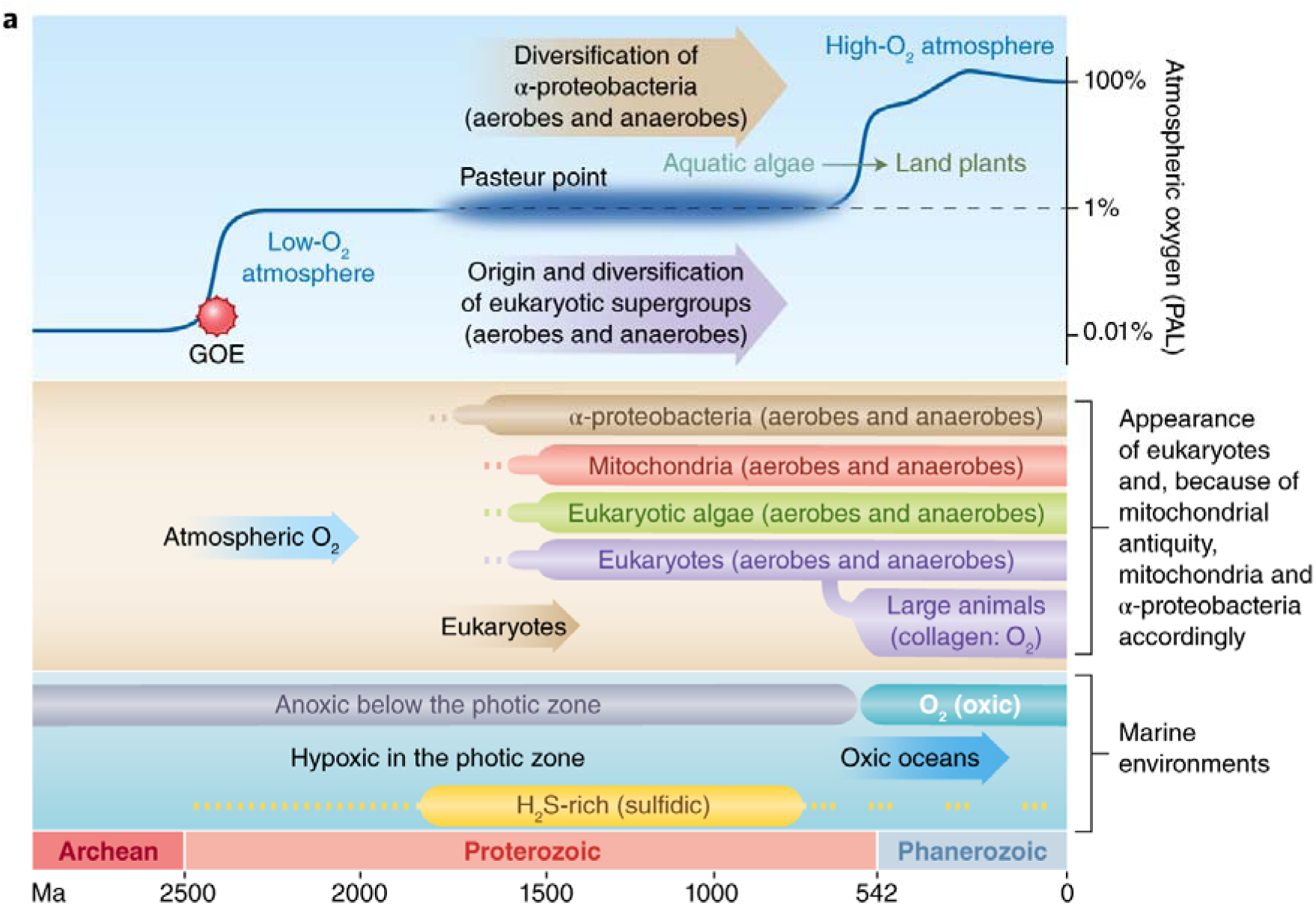
Overview over the changes in Earth’s biochemistry and the rise and diversification of major groups with respect to oxygen concentration. After the great oxidation event (GOE) about 2.45 billion years ago, oxygen concentrations likely fluctuated somewhere around the Pasteur point, as indicated by the cloudy line, with details uncertain and a matter of active debate. What is more certain is the rise of oxygen concentration to extant levels due to a streptophyte alga conquering land some 500 million years ago and the beginning of massive terrestrial carbon burial. Ma, million years ago.

Current findings have it that the monophyletic origin of land plants, which occurred some 450 million years ago (Wickett et al. 2014; de Vries et al. 2016), boosted O_2_ accumulation to modern levels through massive carbon burial (Lenton et al. 2016; Stolper and Keller 2018). Eukaryotes arose roughly 1.8 billion years ago (Parfrey et al., 2011; Betts et al. 2018), from which it follows that the first 1.3 billion years of eukaryote evolution took place in low oxygen conditions (Müller et al. 2012) at atmospheric and marine O_2_ levels comprising only a fraction — 0.001 to 10% — of today’s O_2_ levels (Lyons et al. 2014; Lenton et al. 2016; Stolper and Keller 2018). Because eukaryotes arose and diversified over a billion years before atmospheric O_2_ reached the current value of 21% [v/v], it is hardly surprising that all major lineages (or supergroups) of eukaryotes possess enzymes of anaerobic energy metabolism (Fig. 2). In diverse eukaryotic lineages, these enzymes afford redox balance during ATP synthesis in mitochondria, anaerobic mitochondria (Tielens et al., 2002), hydrogenosomes (Lindmark and Müller 1973; Boxma et al., 2004), and the cytosol (Martin and Müller 1998) without requiring the presence of O_2_ as the terminal acceptor (Müller et al. 2012; Martin and Mentel 2008).

**Fig. 2.**
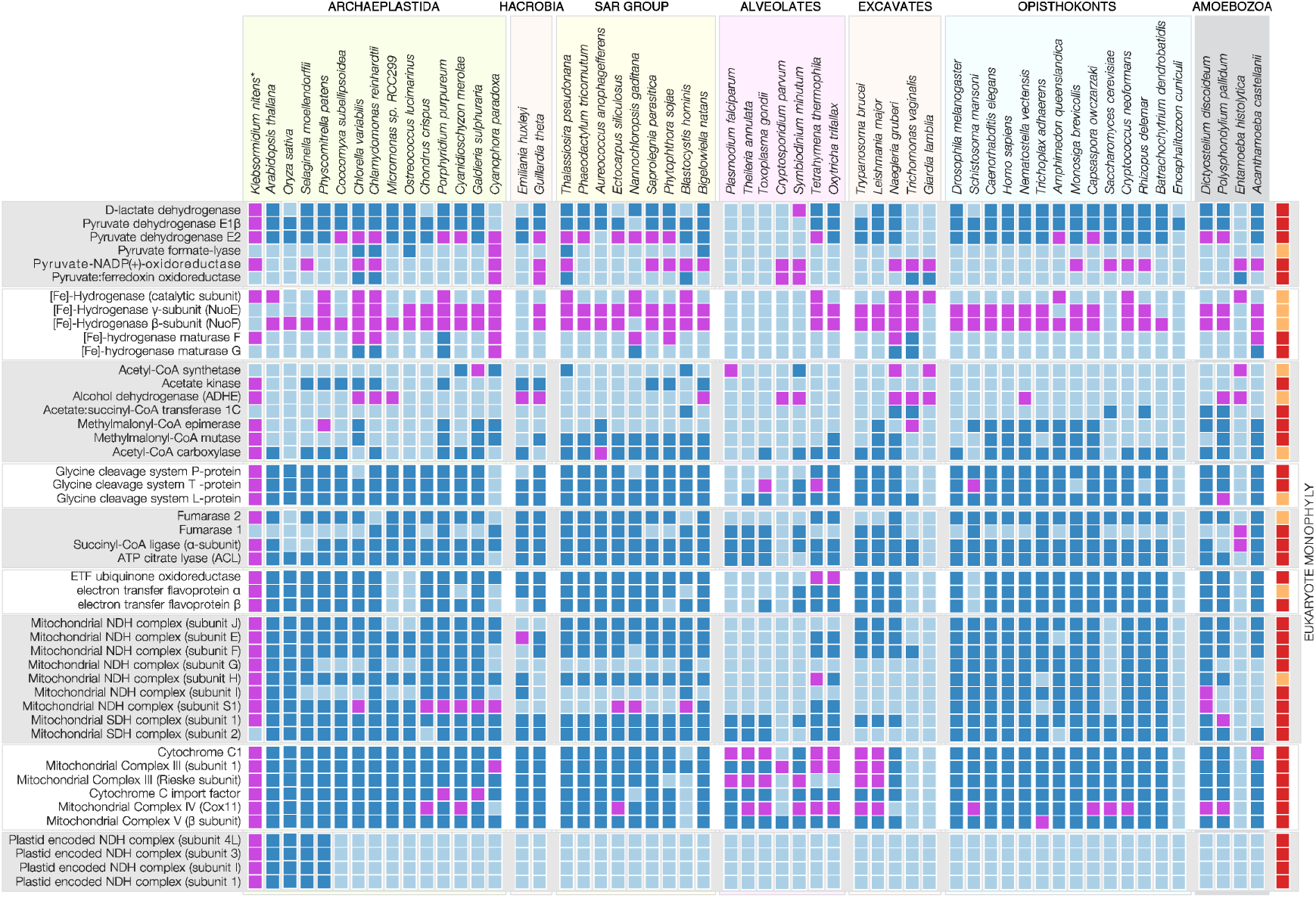
Presence-absence pattern of enzymes associated with anaerobic metabolism across the eukaryotic tree of life. The presence of each enzyme in eukaryotes scored as a dark blue square. An additional BLAST-based search (at least 30% identity and e-value of less than 10^-7^) identifies additional homologs (shown in magenta) that are not represented in the eukaryote-prokaryote clusters (EPCs) from Ku et al. (2015) that is based e.g. on 40% global sequence identity for eukaryote proteins. Enzymes of anaerobic metabolism are present among all eukaryotic supergroups recognized, including all groups of algae, that is those carrying plastids of primary (e.g. *Chlamydomonas reinhardtii, Cyanaphora paradoxa, Volvox carteri*) and secondary origin (e.g. *Bigelowiella natans*). For the enzymes that are identified as EPCs, phylogenetic trees (see supplementary information) indicate that 36/43 (80%) of the genes show a single origin that traces to the eukaryotic common ancestor. Eukaryote monophyly as observed in phylogenetic trees constructed from protein sequences present in each cluster is shown with a dark red (far right column). For an extended PAP including prokaryotes, see supplementary figure 1.

The enzymatic backbone of redox balance in anaerobic energy metabolism in unicellular eukaryotes are pyruvate:ferredoxin oxidoreductase (PFO) and [Fe-Fe] hydrogenase ([Fe]-HYD), which were first described for eukaryotes in studies of carbon and energy metabolism in trichomonad hydrogenosomes (Lindmark and Müller, 1973). The ecophysiological function of these enzymes, together with a larger set of proteins widely distributed across eukaryotes (Fig. 2), is generally interpreted as affording growth without oxygen. Hence, they are typically designated as enzymes of anaerobic metabolism. Like the pyruvate dehydrogenase complex of human or yeast mitochondria, PFO performs oxidative decarboxylation of pyruvate, generating acetyl-CoA and transferring electrons to the 4Fe4S cluster of the one electron carrier ferredoxin (Fd). To maintain redox balance from growth substrate oxidation, reduced Fd (Fd_red_) is re-oxidized by [Fe]-HYD, which donates the electrons to protons, generating H_2_ gas that leaves the cell as a waste product. Fd_red_ generated by PFO is a low potential one electron carrier (*E*_0_’ ca. –400 to –500 mV) that can readily transfer a single electron to O_2_ generating the superoxide radical, O_2_^−^., and reactive oxygen species (ROS). ROS are potent cytotoxins, a reason why organisms that employ the soluble PFO-Fd-[Fe]-HYD electron transport chain avoid high O_2_ environments. In addition, PFO and [Fe]-HYD are irreversibly inactivated by O_2_. Accordingly, eukaryotes that employ PFO and [Fe]-HYD in energy metabolism typically inhabit low oxygen environments, with their possession of these enzymes being interpreted as niche specialization (Martin and Müller 1998, Hug et al., 2010, Stairs et al., 2015).

However, PFO, [Fe]-HYD, and a larger suite of enzymes associated with anaerobic energy metabolism are also present in algae (Ginger et al. 2010; Kruse and Hankamer 2010; Müller et al., 2012; Atteia et al. 2013), phototrophic eukaryotes with plastids that generate O_2_. Their presence in algae is known to enable facultative anaerobic growth in low oxygen environments (Atteia et al. 2013; Müller et al., 2012), and their expression is observed to be upregulated in response to anoxia in algae (Hemschemeier and Happe 2005; Nguyen et al. 2008), in the same way that fermentation enzymes are hypoxia-induced in higher plants (Loreti et al., 2018). However, the expression in O_2_-producing algae of enzymes associated with anaerobic redox balance has not been studied under normoxic conditions. Here, we investigated gene expression data from eukaryotic algae grown at ambient O_2_ levels (21% v/v) to better understand the physiology, function and evolutionary persistence of Fd-dependent enzymes for one electron based redox balance in algae.

### Distribution of enzymes for anaerobic metabolism in eukaryotes

The distribution of 47 genes for enzymes involved in anaerobic energy metabolism (Müller et al. 2012) in 56 eukaryotes spanning the diversity of known lineages is summarized in Fig. 2. The enzymes are widely distributed across diverse eukaryotic lineages, although missing in some, consistent with a standard process of ecological specialization to aerobic and anaerobic habitats entailing the process of differential loss (Ku et al. 2015). Some enzymes of one electron based redox balance have undergone lineage specific functional specialization that entail altered functional constraints in the protein. For example, [Fe]-HYD has lost its H_2_ producing enzymatic activity in several eukaryotic lineages and has assumed different functions. The [Fe]-HYD homologues IOP1/NAR1 repress the hypoxia inducible factor-1α subunit (HIF1-α) in humans (Huang et al. 2007), and furthermore possess conserved functions in cytosolic FeS cluster assembly in human and yeast (Song and Lee 2011; Seki et al. 2013). Prokaryotes employ O_2_-labile FeS clusters for O_2_-sensing and signaling (Barth et al. 2018). In land plants, the [Fe]-HYD homologue has relinquished enzymatic activity to become the oxygen sensor *GOLLUM* (Mondy et al. 2014).

Prokaryotic [Fe]-HYD enzymes can be trimeric (Schut and Adams, 2009), with 24 and 51 kDa subunits associated with the catalytic 64 kDa subunit, which contains the H_2_ producing site, the H cluster. The 24 and 51 kDa subunits allow the enzyme to accept electrons simultaneously from both NADH and Fd via electron confurcation (Schut and Adams, 2009), affording redox balance for both Fd and NADH pools. Some eukaryotic [Fe]-HYD enzymes, including the one from *Trichomonas* hydrogenosomes, also possess the 24 and 51 kDa subunits (Hrdy et al., 2004), which are related to mitochondrial complex I subunits. They are thought to allow the eukaryotes in question to perform electron confurcation, facilitating redox balance via NADH-dependent H_2_ production (Müller et al., 2012), which would be thermodynamically unfavorable in the absence of Fd_red_ (Schut and Adams 2009; Buckel and Thauer 2018).

Intermediate states in the evolutionary transition from Fd-dependent, one-electron based redox balance to NADH dependent redox balance are observed. In various eukaryotic lineages, PFO has become fused to an FAD-FMN-NAD binding domain that converts the ancestrally Fd-dependent enzyme (one electron transport) into an NAD(P)^+^-dependent enzyme that transfers hydride (two electron transport) to generate NADPH. This fusion, called PNO (Rotte et al., 2001), is now known to be widespread among eukaryotes (Fig. 2) (Müller et al., 2012; Stairs et al., 2015), and represents an evolutionary intermediate in the transition from Fd-dependent to NADH dependent redox balance, in that electrons from the FeS clusters of the PFO domain are channeled directly to NAD(P)H, bypassing the generation of soluble Fd_red_.

### Algae express enzymes for anaerobic metabolism at ambient O_2_

The presence of the genes in representatives of the major algal groups (Fig. 2) raises the question of whether and when they are expressed. This is important, because genes for anaerobic energy metabolism have been retained in some eukaryotes with strictly O_2_-dependent energy metabolism (Bexkens et al., 2018). To determine whether enzymes of anaerobic redox balance are expressed independent of anaerobic culturing conditions, we generated transcriptome data for several algal lineages with sequenced genomes: the red alga *Porphyridum purpureum*, the glaucophyte *Cyanophora paradoxa*, the chlorarachniophyte *Bigelowiella natans* with a plastid of secondary green origin, and the cryptophyte *Guillardia theta* with a plastid of secondary red origin. All algae were grown under the same culturing conditions and at ambient O_2_ levels of 21% [v/v]. In all algae, including algae with secondary plastids (Fig. 3a), we were able to detect the expression of at least a subset of the corresponding genes. It is well known that other algae such as *Vitrella brassicaformis* and *Chromera velia* encode a set of anaerobic enzymes that is as complete as that of *C. reinhardtii* (Stairs et al. 2015). We therefore screened available transcriptome data for aerobically grown *Chromera velia* (Woehle et al. 2011; Woo et al. 2015), *Volvox carteri* (Matt et al. 2018), *Chlorella variabilis* (Rowe et al. 2014; Cecchin et al. 2018) and *Thalassiosira pseudonana* (Maheswari et al. 2005) and *Klebsormidium nitens* (de Vries et al. 2018), and find that e.g. the chlorophyte *C. variabilis* and the chromerid *C. velia* (carrying a secondary plastid of red algal origin), express PFL, PNO, HydA/F/G and ADHE in the same way as *C. reinhardtii* for which we generated RNA-Seq data (Fig. 3a).

**Fig 3.**
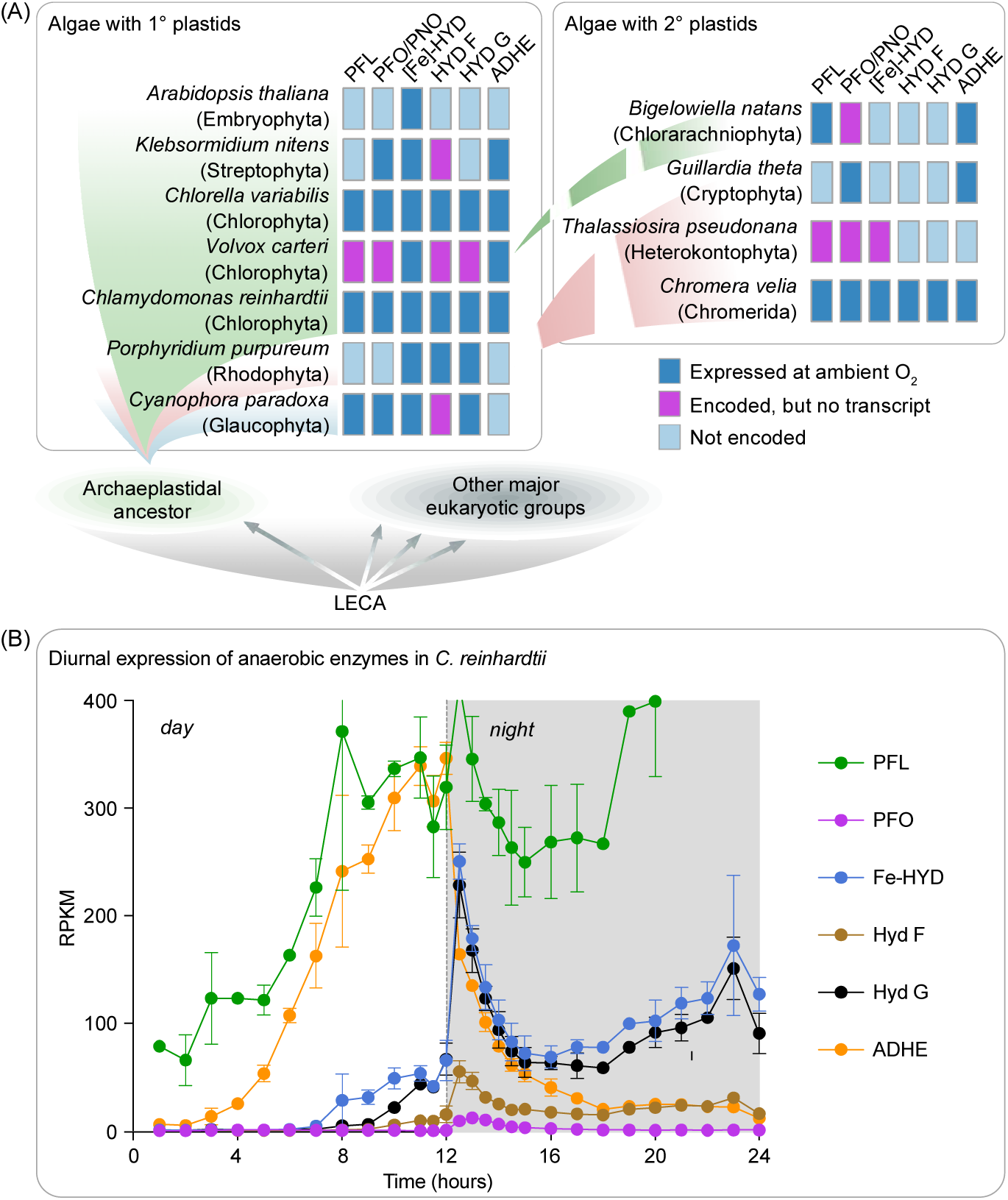
Expression of enzymes of anaerobic metabolism under aerobic conditions in algae and in a diurnal manner in *Chlamydomonas reinhardtii*. (a) Representative algae of all major groups, both with plastids of primary (Glaucophyta, Rhodophyta and Chlorophyta) and secondary origin (Cryptophyta and Chlorarachniophyta), are known to encode enzymes associated with anaerobic metabolism (Fig. 2). We found that those genes for enzymes of anaerobic redox balance are not only encoded in the algae, but are also expressed under ambient O_2_ concentrations and independent of anaerobic growth conditions (except for *T. pseudonana*). (b) High-resolution RNA-Seq data of the chlorophyte alga *C. reinhardtii* shows that the enzymes are mostly expressed in a diurnal manner under aerobic growth conditions. PFL is again seen to be expressed at high levels throughout (to the extent of a house-keeping gene), but with a peak early on during the dark phase that matches that of the other genes in question.

The high-resolution RNA-Seq data available for *C. reinhardtii* (Zones et al. 2015) provided detailed insights into expression of enzymes for redox balance over the time course of 24h. *Chlamydomonas* is among the algae that has preserved the most complete repertoire of O_2_-sensitive enzymes involved in redox balance among eukaryotes studied so far (Fig. 2); it expresses them in the presence of 21% oxygen and in a diurnal fashion (Fig. 3b). PFL is found to be constantly expressed, but more so during the dark phase and in particular towards the end of the night (consistent with our RNA-Seq data). The same pattern is observed for its activating enzyme, although at much lower levels, similar to what is observed in prokaryotes (Crain and Broderick 2014). Other genes in question, including both genes for the [Fe]-HYD catalytic subunit, HydA1 and HydA2, are upregulated with the onset of night (Fig. 3b). Importantly, this induction is observed independent of anaerobic culturing conditions, the standard method employed to induce [Fe]-HYD expression, typically in the context of biohydrogen applications (Vignais et al. 2001; Forestier et al. 2003; Mus et al. 2007; Hemschemeier et al. 2008). The *Chlamydomonas* relatives *Chlorella* and *Volvox* display similar induction of enzymes involved in H_2_ production and dark fermentation (Meuser 2011; Cornish et al. 2015), hence anaerobiosis-independent expression is conserved and *Chlamydomonas* is the rule, not an exception.

The main finding from Figure 3 is that the expression of the enzymes for anaerobic redox balance in eukaryotes does not correspond to any form of adaptation to anaerobic niches, as ambient O_2_ does not change during the 24-hour cycle. Instead, their expression in *C. reinhardtii* corresponds to the onset and end of illumination, where electron flux to and from the photosynthetic electron transport chain undergoes transient changes. In *Chlamydomonas*, PFO and [Fe]-HYD are localized in the plastid (Hemschemeier et al. 2008), not the mitochondrion or the cytosol, they help to buffer electron flow into and out of the thylakoid membrane. This function does not preclude the existence of other functions under other conditions. For example, the same genes are expressed in *C. reinhardtii* during anaerobiosis (Hemschemeier and Happe 2005; Nguyen et al. 2008). Yet for most of the algae surveyed in Figure 3a, extended anaerobic growth phases are unknown, and the main habitat is the photic zone, where daily diurnal light conditions are encountered.

Some might view *C. reinhardtii* as an extreme case among algae, as it appears to mimic true anaerobic protists such as *Trichomonas* or soil-dwelling anaerobic bacteria when experiencing hypoxia. But *Chlamydomonas* can only endure anaerobic conditions for a limited amount of time, not thrive under them. Sustained hydrogen production by *Chlamydomonas* for an elongated period of time is only feasible when PSII provides the main source of reducing power (Scoma et al. 2014). This underscores our point that photosynthetic redox balance and one electron based redox balance conferred by the soluble PFO-Fd-[Fe]-HYD electron transport chain operate independent of anaerobiosis, because PSII activity entails O_2_ production. Finally, *C. reinhardtii* is not the only alga encoding such a complete set of anaerobic enzymes (Atteia et al., 2013; Stairs et al. 2015), but the only one that has been extensively studied in this respect.

## Discussion

The retention and anaerobiosis-independent expression of Fd-dependent enzymes in algae, together with their localization to plastids in cases studied to date, indicates that the enzymes have been retained during algal evolution as the result of selection for redox balance in cells with one electron transport. In terms of gene distribution (Fig. 2) and phylogeny (Supplementary Information Fig 1), the enzymes of anaerobic energy metabolism in eukaryotes trace to the eukaryote common ancestor (Ginger et al. 2010; Atteia et al. 2013) (Fig. 2), hence the archaeplastidan founder lineage that acquired the cyanobacterial ancestor of plastids already possessed them.

Eukaryotic enzymes involved in Fd-based redox balance have been the subject of many evolutionary investigations. There are two alternative hypotheses to account for their presence in eukaryotes. One has it that the Fd-dependent enzymes were present in the eukaryote common ancestor, which was a facultative anaerobe that was able to survive with or without O_2_ as terminal acceptor, and were involved in its energy metabolism and redox balance (Martin Müller 1998, Tielens et al 2002; Müller et al., 2012; Degli Esposti 2014). The alternative lateral gene transfer (LGT) hypothesis has it that the ancestral eukaryote was a strict aerobe, unable to survive under anaerobic conditions, the presence of the Fd-dependent enzymes in eukaryotes resulting from multiple lateral gene transfers during eukaryote evolution to confer the ability to colonize anaerobic niches (Stairs et al. 2011; Stairs et al. 2015; Eme et al. 2017). Directly at odds with the LGT theory is the observation that the archaeplastidal ancestor, whose PFL and PFL-activating enzyme are of monophyletic origin (Stairs et al. 2011), did not adapt to an anaerobic niche, rather it acquired a cyanobacterial endosymbiont that became an O_2_-producing plastid. The archaeplastidal lineage diversified into three main groups, representatives of which have retained the enzymes (Atteia et al. 2013; Stairs et al. 2015) (Fig. 2).

Though various formulations of the LGT hypothesis for enzymes of anaerobic redox energy metabolism in eukaryotes differ with respect to the number, nature and direction of LGTs (Stairs et al. 2015), the underlying evolutionary rationale of the LGT hypothesis has remained constant: the lateral acquisition of Fd-dependent enzymes supposedly allowed eukaryotes to colonize oxygen poor niches (Leger et al. 2018). Notwithstanding the circumstance that the majority of eukaryote evolution occurred in oxygen poor environments (Martin and Mentel, 2008; Müller et al., 2012; Lyons et al., 2014; Lenton et al. 2016) (Fig. 1), the diurnal expression of Fd-dependent enzymes in algae at 21% [v/v] O_2_ (Fig. 3) and independent of anaerobic growth conditions is incompatible with the view that the presence of these genes has anything to do with lateral acquisitions for adaptation to anaerobiosis. Rather, the data indicate that the genes for Fd-dependent redox balance were present in the eukaryote common ancestor, lost in some lineages during specialization to permanently oxic habitats (Fig. 4), and retained in lineages that did not undergo the irreversible adaptation to complete O_2_ dependence and NADH dependent redox balance (Fig. 1).

**Fig. 4.**
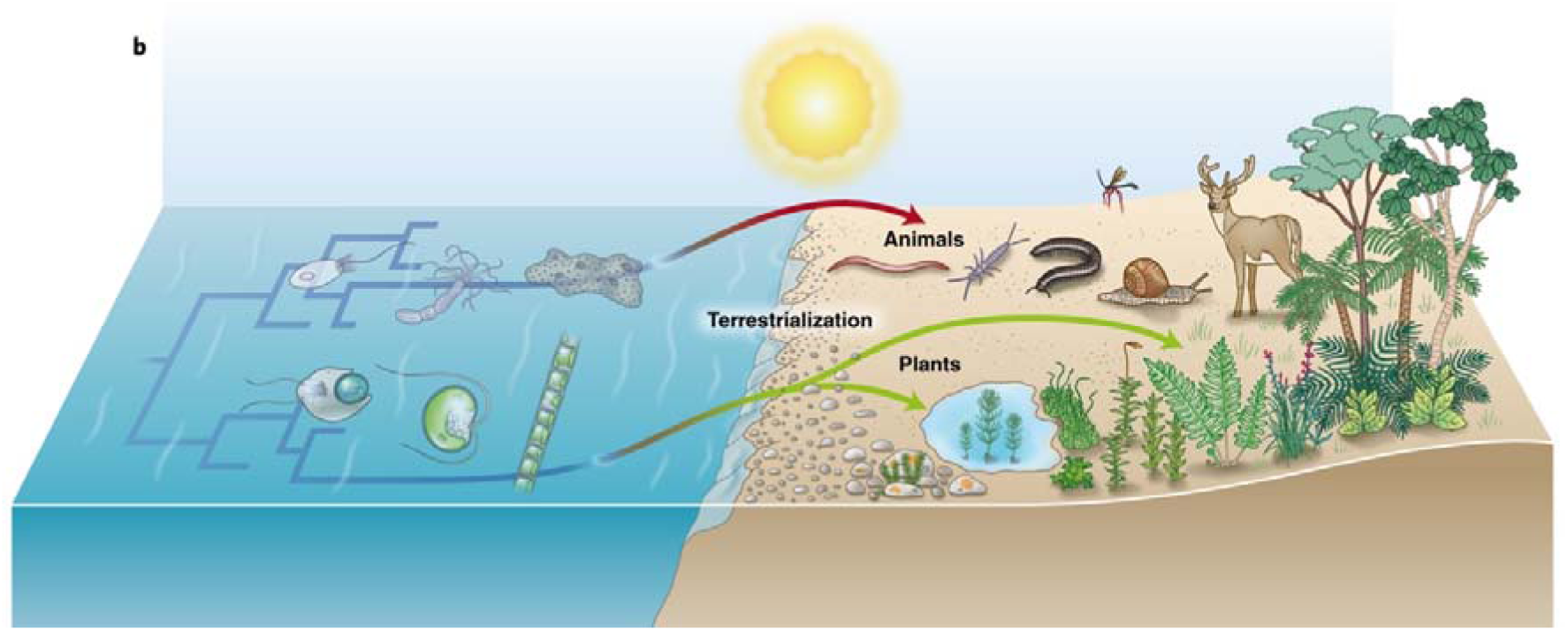
Animal and plant terrestrialization. The major eukaryotic groups emerged in aquatic environments, including the Archaeplastida that stem from the endosymbiotic integration of a cyanobacterium into a heterotrophic protist. The conquer of land by a streptophyte alga preceded that of animals. The rise in oxygen concentration fostered the evolution of macroscopic life adapted to high oxygen concentrations, which includes NADH-(two electron) based redox balance.

Evolutionary responses to redox balance in eukaryotes can include recompartmentalization of pathways (Martin 2010) to the cytosol, to plastids (Schnarrenberger et al., 1995), to glycosomes (Opperdoes and Borst 1977), or to mitochondria (Rio Bartulos et al., 2018). Based upon their presence in the eukaryote common ancestor and their current localization within plastids in algae studied to date, the Fd-dependent enzymes PFO and [Fe]-HYD were recompartmentalized to the plastid during algal evolution. In the plastid they assumed essential roles in light-dependent redox balance in an organelle that, upon contact with light, has no choice but to commence photosystem I dependent Fd reduction, rapidly depleting the available Fd_ox_ pool. In land plants, Fd_red_ is mainly reoxidized by ferredoxin:NADP^+^ oxidoreductase (FNR), NADPH being reoxidized in turn by NADP^+^-dependent glyceraldehyde 3-phosphate dehydrogenase (GAPDH) (Cerff 1982) in the Calvin cycle. In aquatic environments, CO_2_ is more limiting than in air, for which reason algae have evolved diverse CO_2_ concentrating mechanisms (Giordano et al., 2005). Algae thus require a means in addition to CO_2_ fixation for redox balance at the onset of illumination, Fd-dependent enzymes of anaerobic energy metabolism fulfill that role. That functional aspect, redox balance in the plastid rather than anaerobiosis, accounts for diurnal expression and retention of enzymes for anaerobic redox balance among many independent algal lineages (Fig 1). The expression of ferredoxin dependent enzymes thus enables redox balance in the presence of O_2_ in plastids and in the absence of O_2_ as it occurs in *C. reinhardtii* (Hemschemeier and Happe 2005) and many lineages of anaerobic protists that arose and diversified before the origin of plastids (Martin and Müller 998; Müller et al., 2012).

The transition to life on land ∼450 million years ago marked the advent of life in very high O_2_ conditions (Lenton et al. 2016; Stolper and Keller 2018). Plants were the first major colonizers of land (Nishiyama et al. 2018). Massive carbon burial by land plants precipitated the high O_2_ environment into which the first land animals followed (Fig. 4). The colonization of land was, physiologically, an adaptation to high O_2_ air. That adaptation to high O_2_ witnessed the loss of Fd-dependent redox balance independently in both the land plant and land animal lineages (Fig. 4) in response to the O_2_ sensitivity of FeS clusters in PFO and [Fe]-HYD and in response to the ROS generating potential PFO of Fd_red_. Once on land, both the plant and animal lineages were subsequently confronted again with hypoxic environments in adaptations to aquatic environments. The corresponding adaptations did not, however, involve gene acquisitions via LGT, merely novel expression regulation for NADH dependent enzymes involved in redox balance during hypoxic response. In plants, these responses include mainly ethanol fermentations in waterlogged roots (Perata and Alpi, 1993, Licausi et al., 2010; Loreti et al 2018). In animals, the evolutionary responses include various pathways regulated by the hypoxia induced factor HIF (Semenza 2001; Kim et al. 2006), and secondary adaptations to the aquatic lifestyle among various vertebrates (Hochachka and Somero, 2002; Darveau et al., 2002). In addition, many marine and soil dwelling invertebrates independently evolved their own specialized strategies for redox balance (Müller et al. 2012), from opine accumulation in mussels (Grieshaber and Völkel 1998) to rhodoquinone dependent short chain fatty acid excretion in worms (Komuniecki et al., 1998). The land plant and land animal anaerobiosis adaptation pathways are, however, always NADH dependent.

The retention of the chloroplast encoded NADH dehydrogenase complex (cpNDH) specifically in the land plant lineage (Fig. 1) is noteworthy. The functional cpNDH complex is localized close to complex I in thylakoids, both in the cyanobacterium *Synechocystis* (Mi et al. 1995) as well as in land plants, where it supports the cyclic flow of electrons essential for PSI to properly perform photosynthesis (Munekage et al. 2004; Yamori et al. 2011). Among genes in plastid DNA, the cpNDH genes have undergone the highest number of independent losses (Martin et al. 2002). Their retention in the plastid was likely a prerequisite for the transition to life on land (de Vries et al. 2016; Nishiyama et al. 2018), because they have been retained by the plastid in all land plant lineages, indicating a strong functional constraint for maintaining redox balance in the organelle (Allen 2015). Land plants have recruited a cytosolic NADH dependent GAPDH (Petersen et al. 2003) and a cytosolic malate dehydrogenase (Selinski et al. 2014) to plastids for NADH-based redox balance. Even the origin of photorespiration, a process central to NAD(P)H dependent redox balance, can be understood as an evolutionary response to high O_2_ in the transition to life on land (Hanawa et al, 2017). Land plant thylakoids cannot, however, relinquish Fd-dependent one electron transport altogether, because the structure and function of photosystem I strictly require a steady flow of single electrons from the FeS clusters of PSI to generate soluble Fd_red_, the stromal levels of which are monitored in some photosynthetic lineages by the flavodiiron (FLV) proteins (Gerotto et al. 2016). Our findings indicate that in the plant and animal lineages, terrestrialization entailed an irreversible physiological transition away from one electron based Fd-dependent redox balance towards NAD(P)H-dependent redox balance involving two electron transfers. The underlying evolutionary mechanisms were gene expression changes, enzyme recompartmentalization and gene loss in adaptation to high O_2_ levels. Algae retained the Fd-dependent pathway for Fd-dependent, one-electron based redox balance in plastids, not for anaerobic growth.

## Material and Methods

### Identification of homologous proteins

As part of a larger study (Ku et al., 2015) sequences from 55 eukaryotes and 1,981 prokaryotes (1,847 bacteria and 134 archaea) were clustered into protein families in order to identify eukaryotic proteins with prokaryotic homologs. This approach resulted in 2,585 disjunct clusters that contain at least two eukaryotes and no less than five prokaryotes. Within these 2,585 eukaryote-prokaryote clusters (EPCs) using existing annotations we identified 42 clusters containing proteins involved in anaerobic energy metabolism, which were relevant for the current analysis (Supplementary Table 1). Phylogenetic trees and results from the tests on eukaryote monophyly were taken directly from the analysis performed in Ku et al., 2015 (shown in Supplementary Table 1). For proteins that did not have an EPC in Ku et al., 2015 the same dataset was used to perform a BLAST search and only hits with an identity of greater than 30% and an e-value less than 10^-10^ were considered and provided in supplementary table S2. All the sequences that were used to identify the EPCs and perform the BLAST search are provided in Supplementary File 1 along with the BLAST hits.

### Cultivation of algae, RNA isolation and transcriptomics

All algae were grown in their respective media (see SAG Göttingen or ncma.bigelow.org) in aerated flasks under controlled conditions at 22°C and illuminated with 50µE under a 12/12h day-night cycle. RNA was isolated from cells growing in the exponential phase and at 6h into the day and 6h into the night. RNA was isolated using either Trizol™ reagent (Thermo Fisher, Cat. No.: 15596018) or the Spectrum™ Plant Total RNA Kit (Sigma Aldrich, Cat. No.: STRN50) according to the manufacturer’s protocols. Then, samples were DNase treated (DNase I, RNase free, Thermo Fisher, Cat. No.: EN0525) and RNA-Seq was performed at the Beijing Genome Institute (BGI, Hong Kong) using an Illumina HiSeq2000 resulting in 150bp paired-end reads. For each sample, three individual runs were performed and pooled. Raw reads were subjected to several cleaning steps. First, adapter sequences as well as reads containing more than 5% of unknown nucleotides or more than 20% of nucleotides with quality scores less than 10 were removed. Further, reads were processed using trimmomatic (version 0.35) (Bolger et al. 2014) by removing the first 10 nucleotides as well as reads which showed a quality score below 15. Additionally, poly-A/T tails ≥ 5 nt were removed using prinseq-lite (version 0.20.4) (Schmieder & Edwards 2011). Finally, only reads with a minimum length of 25 nt were retained. Trimmed reads were assembled using Trinity (version 2.2.0) (Grabherr et al. 2011) and resulting contigs were filtered for a minimum length of 300 nt using an in-house perl script. Subsequently, open reading frames (ORFs) were identified using Transdecoder (version 3.0.1) (https://github.com/TransDecoder/TransDecoder/wiki). These ORFs were used for a BLAST with an identity cut off of 30% against the genome of the respective organisms to verify their presence in the genome and to remove possible contaminations. Transcriptomes are available at [*accession numbers will be made available once the manuscript has been accepted*].

## Acknowledgments

We gratefully acknowledge the funding of this work by the DFG (to SBG; 267205415 – SFB 1208), the ERC (WFM; 666053), and the VolkswagenStiftung to SBG and WFM (both Life).

## Supplementary files

**Supplementary Figure 1**: Presence-absence pattern of enzymes associated with anaerobic metabolism across and prokaryotes.

**Supplementary Table 1**: Annotations of the functions of the eukaryote prokaryote clusters from Ku et al., 2015 and eukaryote monophyly in EPC trees.

**Supplementary Table 2:** BLAST output for the sequences without EPCs in Ku et al., 2015.

**Supplementary Table 3:** FASTA headers of all the sequences used in the study along with their aliases.

**Supplementary Table 4:** BLAST output of the expressed contigs against their respective genomes.

**Supplementary File 1**: FASTA sequences of the proteins screened for in the expression data.

**Supplementary File 2**: Extracted Newick trees (from Ku et al. 2015) for clusters containing analyzed enzymes that are part of figure 2. Columns: (i) enzyme name; (ii) enzyme name abbreviation; (iii) cluster number as referenced in Ku et al. 2015; (iv) tree in Newick format.

**Supplementary File 3:** FASTA sequences of the expressed contigs that mapped to the respective genomes.

