## Supplementary File 3 for "Adaptation to life on land at 21% O_2_ via transition from ferredoxin- to NADH-dependent redox balance"

>Cr_TRINITY_DN14203_c6_g1::TRINITY_DN14203_c6_g1_i1::g.157573::m.157573
MLTPLSYPIINTVSSSLPALHAMSQMLLEKTMRRGLATVSAAASSAVGRPIPMAVRPSLTLGNVRGMRLGVPELAFRSPMRSMAAASAAAEALPVAPSHSCAADPDKHPHLPDPRPKPAVDAGINVQKYVQDNYTAYAGNSSFLAGPTDNTKKLWSELEKMIATEIEKGVMDVDPSKPSTITAFPPGYIDKDLETVVGLQTDAPLKRAIKPLGGVNMVKAALESYGYTPDPEVARLYSTVRKTHNSGVFDAYTDEMRAARKSGILSGLPDGYGRGRIIGDYRRVALYGVDALIKAKKTDLKHNLLGVMDEEKIRLREEVNEQIRALSELKEMGAAYGFDLSRPAANSREAVQWLYFGYLGAVKEQDGAAMSLGRIDAFLDTYFERDLKAGTITEAEVQELIDHFVMKLRIVRQLRTPEYNALFAGDPTWVTCVLGGTDASGKAMVTKTSFRLLNTLYNLGPAPEPNLTVLWNDNLPAPFKEFCAKVSLDTSSIQYESDNLMSKLFGSDYSIACCVSAMRVGKDMQYFGARANLPKLLLYTLNGGRDEVSGDQVGPKFAPVRSPTAPLDYEEVKAKIEDGMEWLASMYANTMNIIHYMHDKYDYERLQMALHDTHVRRLLAFGISGLSVVTDSLSAIKYAQVTPVIDERGLMTDFKVEGSFPKYGNDDDRVDEIAEWVVSTFSSKLAKQHTYRNSVPTLSVLTITSNVVYGKKTGSTPDGRKKGEPFAPGANPLHGRDAHGALASLNSVAKLPYTMCLDGISNTFSLIPQVLGRGGEHERATNLASILDGYFANGGHHINVNVLNRSMLMDAVEHPEKYPNLTIRVSGYAVHFARLTREQQLEVIARTFHDTM*
>Cr_TRINITY_DN14437_c5_g2::TRINITY_DN14437_c5_g2_i1::g.6156::m.6156
ITKAYERKGPEVVAKNHSAVDMAVAALKKLDIPASWSSLPTHVVNPNPPAKGNTSRWEFIETVAKPMLALEGDKLPVSVFSPEGFVPPGTTVIEKRAIAAQVPIWKAENCTQCNVCAFVCPHAAIRPALASPADLVGAPATFGTIQAKGPGMGDLKYRIQVSPYDCTGCDLCTHACPDDALQSVPINSVLEVETANWDFAGTLPARTDIMDKATVKGSQLQPPLMEFSGACEGCGETPYVKLLTQLFGDRLIIANATGCSSIWGGSAPSNPYTTNADGYGPAWANSLFEDNAQFGLGIAMGTMQRRKTVRKHVQEVLAVEAEKMPLSFGLRNALTRWDEHFDEPEVANVVAKELQPLLEREKDVHPWIRQLYDERDMLHKASIWIVGGDGWAYDIGFGGLDHVLASGENVNILVLDTEVYSNTGGQRSKATPKSAVTKFAAGGKERPKKDLGAIAMSYGDVYVASTSLHANYGQVTKAMAEAEKYGGVSLVLAYAPCIMHGISSGMCSAIDESKQAVETGYWPLYRYNPAVKEDATHHRFQLDAKKLKGDVEEWLSHENRFQILERKNPEVAHKLHHELDDAVHERFDRMKHMAAGGHMEPGSPPPPAAAAPQAQAPDHGHQHNTKESGCGGH*
>Cr_TRINITY_DN13317_c0_g1::TRINITY_DN13317_c0_g1_i5::g.207267::m.207267
MALGLRAELRAGQAVACARRTNAPAHPAAVVPVLPSRGDKFFNLSQKVPSSQPARGSTIRVAATATDAVPHWKLALEELDKPKDGGRKVLIAQVAPAVRVAIAESFGLAPGAVSPGKLAAGLRALGFDQVFDTLFAADLTIMEEGTELLHRLKEHLEAHPHSDEPLPMFTSCCPGWIAMLEKSYPDLIPYVSSCKSPQMMLAAMVKSYLAEKKGIAPKDMVMVSIMPCTRKQSEADRDWFCVDADPTLRQLDHVITTVELGNIFKERGINLAELPEGEWDNPMGVGSGAGVLFGTTGGVMEAALRTAYELFTGTPLPRLSLSEVRGMDGIKETNITMVPAPGSKFEELLKHRAAARAEAAAHGTPGPLAWDGGAGFTSEDGRGGITLRVAVANGLGNAKKLITKMQAGEAKYDFVEIMACPAGCVGGGGQPRSTDKAITQKRQAALYNLDEKSTLRRSHENPSIRELYDTYLGEPLGHKAHELLHTHYVPGGAEADA*
>Cr_TRINITY_DN13205_c1_g2::TRINITY_DN13205_c1_g2_i1::g.121746::m.121746
ASARLCAGARRAGRVVASPLRPAAACRGVAVKAAAAAAGEDAGAGTSGVGSNIVTSPGIASTTAHGVPRINIGVFGVMNAGKSTLVNALAQQEACIVDSTPGTTADVKTVLLELHALGPAKLLDTAGLDEVGGLGDKKRRKALNTLKECDVAVLVVDTDTAAAAIKSGRLAEALEWESKVMEQAHKYNVSPVLLLNVKSRGLPEAQAASMLEAVAGMLDPSKQIPRMSLDLASTPLHERSTITSAFVKEGAVRSSRYGAPLPGCLPRWSLGRNARLLMVIPMDAETPGGRLLRPQAQVMEEAIRHWATVLSVRLDLDAARGKLGPEACEMERQRFDGVIAMMERNDGPTLVVTDSQAIDVVHPWTLDRSSGRPLVPITTFSIAMAYQQNGGRLDPFVEGLEALETLQDGDRVLISEACNHNRITSACNDIGMVQIPNKLEAALGGKKLQIEHAFGREFPELESGGMDGLKLAIHCGGCMIDAQKMQQRMKDLHEAGVPVTNYGVFFSWAAWPDALRRALEPWGVEPPVGTPATPAAAPATAASGV*
>Cr_TRINITY_DN14295_c2_g2::TRINITY_DN14295_c2_g2_i1::g.199706::m.199706
MSVPLQCNAGRLLAGQRPCGVRARLNRRVCVPVTAHGKASATREYAGDFLPGTTISHAWSVERETHHRYRNPAEWINEAAIHKALETSKADAQDAGRVREILAKAKEKAFVTEHAPVNAESKSEFVQGLTLEECATLINVDSNNVELMNEIFDTALAIKERIYGNRVVLFAPLYIANHCMNTCTYCAFRSANKGMERSILTDDDLREEVAALQRQGHRRILALTGEHPKYTFDNFLHAVNVIASVKTEPEGSIRRINVEIPPLSVSDMRRLKNTDSVGTFVLFQETYHRDTFKVMHPSGPKSDFDFRVLTQDRAMRAGLDDVGIGALFGLYDYRYEVCAMLMHSEHLEREYNAGPHTISVPRMRPADGSELSIAPPYPVNDADFMKLVAVLRIAVPYTGMILSTRESPEMRSALLKCGMSQMSAGSRTDVGAYHKDHTLSTEANLSKLAGQFTLQDERPTNEIVKWLMEEGYVPSWCTACYRQGRTGEDFMNICKAGDIHDFCHPNSLLTLQEYLMDYADPDLRKKGEQVIAREMGPDASEPLSAQSRKRLERKMKQVLEGEHDVYL*
>Cr_TRINITY_DN14556_c0_g1::TRINITY_DN14556_c0_g1_i4::g.197831::m.197831
MMSSSLVSGKRVAVPSAAKPCAAVPLPRVAGRRTAARVVCEAAPSGAAPASPKAEAAAPVAAAPATPHAEVKKERAPATDEALTELKALLKRAQTAQAQYSTYTQEQVDEIFRAAAEAANAARIPLAKMAVEETRMGVAEDKVVKNHFASEFIYNKYKHTKTCGVIEHDPAGGIQKVAEPVGVIAGIVPTTNPTSTAIFKSLLSLKTRNALVLCPHPRAAKSTIAAARIVRDAAVAAGAPPNIISWVETPSLPVSQALMQATEINLILATGGPAMVRAAYSSGNPSLGVGAGNTPALIDETADVAMAVSSILLSKTFDNGVICASEQSVVVVAKAYDAVRTEFVRRGAYFLTEDDKVKVRAGVVVDGKLNPNIVGQSIPKLAALFGIKVPQGTKVLIGEVEKIGPEEALSQEKLCPILAMYKAPDYDAGVKMACDLIMYGGAGHTSVLYTNPLNHAHIQQYQSAVKTVRILINTPASQGAIGDLYNFHLDPSLTLGCGTWGSTSVSTNVGPQHLLNIKTVTARRENMLWFRVPPKIYFKGGCLEVALTDLRGKSRAFIVTDKPLFDMGYADKVTHILDSINVHHQVFYHVTPDPTLACIEAGLKEILEFKPDVIIALGGGSPMDAAKIMWLMYE
>Bn_TRINITY_DN24900_c0_g1::TRINITY_DN24900_c0_g1_i1::g.109868::m.109868 TRINITY_DN24900_c0_g1::TRINITY_DN24900_c0_g1_i1::g.109868  ORF type:complete len:755 (-) TRINITY_DN24900_c0_g1_i1:686-2950(-)
MLQASLRRLAPAASIIAKRMAPLASSALCRMARRNYQVYSGDSSFLEGPTKRTELVWGQCLDLCKEELKNGILDVDNKIPSDVNAFPPGYIDKENEVIVGLQTDKPLKRSIKPYGGLRMVKSALESYGYSLDPDTEAIFRKYRKTHNEGVFDVYTDEMKTARKYKILTGLPDAYGRGRIIGDYRRVAKYGVSHLINQKILDKKNLGGKMDEHSIRLAEEIAEQIKALKELVKMAEGYGFDIMRPAENAKEATQWLYFAYLGAIKQQDGAAMSLGRVDCFLDEYFEKDLANGVLSESDAQEIIDDFVIKLRLVRHLRTPEYNELFAGDPQWITLVIGGLDEKGQPRVTKTSYRILQTLYNLGPGPEPNLTILWSERLPEHFKRFCAQSSIAMSSIQYENDDLMRPRFSDDYGIACCVSAMNIGEQMQFFGARCNLAKLLLYAINQGVDEISGAQVGPRFAPVSEGAGGGVLRYEEVMERYRAAIEWLAELYVNTMNVIHYMHDKYSYESLQMALHDSEVKRFMAFGVAGLSSAADSLSAIKYAKVTPVRDERGIATEFKVEGDFPAYGNNDDRVDEIARDLLDAFSGELKKTPTYRDSEHTLSVLIITSNVVYGKKTGSTPDGRIKGEPFAPGANPMHGRDNSGCINSLKSVAKMSYDSCMDGISNTFTVIPSALGKNEPDRESNLVSLIDGYFQDGAQHLNVNVLNREMLIDAAENPEKYPNLTIRVSGYAVHFNRLTKEQQQDVISRTFHAAM*
>Bn_TRINITY_DN9763_c0_g2::TRINITY_DN9763_c0_g2_i2::g.234271::m.234271 TRINITY_DN9763_c0_g2::TRINITY_DN9763_c0_g2_i2::g.234271  ORF type:complete len:882 (+) TRINITY_DN9763_c0_g2_i2:119-2764(+)
MLLSAKCLRGTFRRTHRIASFRYISTSTEDKLERVKRASEIFATFSQERVDYIFTRIAHEANKQRVPLAQIAHEETGMGLMEDKVIKNGIACELLLDRYRDTKTCGIIERESVHGLLKIANPVGVVCCITPTTNPTSTAIAKSLFCAKTRNGAIFLPHPRALESTAAAVKVCHDAGVAAGAPKGFLECVDEVNREVSDFVMHHDNVNIIMATGGPGMVKASYSTGKPALGVGSGNAPILVDETANLEEACGSIVLGKTFDNGMICAAEQSVVCVESVYDQFKEMLIRRGVTFLEGKERDALGKFLIVNGRVNADIVGQSAQEIARRINVEVPYGTVVLGAEASNVGPQEPFSHEKLCPVLALYKASDFKEGVKLCKELTEYDGIGHTAGIYSRSQERLEEYAQKIPAGRILNNVPTSLAAIGTAFNFNIDFSLTLGVGTKAGSSVSDNVGPMHLLNVRTVATRQEHTEWFKNPPQIYFNRNCLEDALRDVSNKYADGTKDQRAIIITDEVMGSLGYVERVKRCLKDKGFVVSVFDDVHPDPDIMTVRKGVKACEAFRPDLMVCLGGGSPMDAGKYIRVSYEHPELHLEDAASRFIEIRKRTKHFPKLGSKIRKLVCIPTTSGTASEVTPFSVITDDMGMKHPLFDYELTPDIAIIDSSFCDKLPKSLIAHAGVDAITHATEAFVSVAANEFTESHSLTAINLLFQHLPNSYHTGDPQAREMVHHAATLAGLAFSNSYLGITHSLSHKVGAVFHLPHGLTNAVLMPYVIRFNASERPTRMGIYPGYDHPIAKQRYAAIATHLGLKGKTDDELCEAYCQAILGLMDNLEMPRTFEKCGVNEKAYFEQIDDMALAAFDDQCTPANPRFPLVSELKQILTDSYHG*
>Cp_TRINITY_DN25614_c11_g4::TRINITY_DN25614_c11_g4_i1::g.110726::m.110726 TRINITY_DN25614_c11_g4::TRINITY_DN25614_c11_g4_i1::g.110726  ORF type:internal len:1800 (-) TRINITY_DN25614_c11_g4_i1:2-5398(-)
PKSAQRFSAKLFSRLASIAHSFVQRTPLTRAAAWDRGLSRMACPMSAAAPAAKAPTFKSLDGNEAAASVAYAFSDIAFIYPITPSSPMAEHADEWSAMGKPNLFGQVVQVKEMESEAGAAGALHGALAAGALGTTFTASQGLLLMIPNMYKIAGELMPCVMHVAARSLAGQALCIFGDHSDVMAVRQTGWALLSAATVQEAYDLAIVSHLSTLKSRVPFVHFFDGFRTSHEVQKIDATPLSHFKTFMDMKAIEDHRGRALNPAHPHMRGTAQGSDVFFQNVEAANQFYDKVPAIVEQTMNEVASVTGRPFKLVDYYGAPDAELVIVSMGSSCPVIEETIDALKGQKIGLIKIRLFRPWPHAQFLETLPKTTKRICVLDRTKEAGAQGEPIYLDVCTSLHEGEVTGKLVVGGRYGLGSKEFTPSMVKAIYDNLALPKPKNHFTVGIVDDVTHTNLPLGPPMRTVPEGTVQCQFWGMGSDGTVGANKDAVKIIGDNSELFTQAYFSYDAHKSGGLTVSHLRFGPKPITSTYLVTEADYVAVHLESYVTKYNFLATLRPGGAVVLNTVWTAAELEKKLPASVKRQLHELRARLYIINARAISDSVGLGKRINMVMQTVFFYLSGVLPFERAIALLKKSIEKTYGKKGPEIVAMNHRCVDAAVAHLVTVPIPDEWKTAQDDTRRRAPGPVATDFVKNVVKPMLAMEGDSLPVSAFVPGGVVPPGTTQYEKRAIASKVPTWKADKCSQCNYCAFVCPHAAIRPVLVTEEELEGGAPEAFETRKAKGSGEITNFKYRVQVSPYDCTGCELCIHACPDEALEAAEINDILDTEAANWDYFTTVPPRGELFDRNTVKGSQFQQPLLEFSGACEGCGETPYVKLLTQLFGERMIIANATGCSSIWSGSAPVNPYTTNKDGRGPAWANSLFEDNAQFGYGIATGVIQRRAGLRKVIDDVATTPEGQPGFVPMSSELRSAMKAWLDVWQDADKCESASKRLETVLGAECRRPGSSAMLREIYKQKDLLTKASIWIVGGDGWAYDIGYGGLDHVLASGENVNILVMDTEMYSNTGGQQSKSTPMASVAQFAMSGKKQNKKDLGILAMQYGSVYVASVALGANFSQVTRAMVEAERYPGVSLIIAYAPCMLHGIREGMSYALDESKMAVDTGYWTLYRFDPRIKVDAEHTQFQLDSKKIKAELTTFLKHENRFGILARSCPSEAKKLQVNLQEFVIARHEKYKHLAETDAERKANLAAQFAKLAGGLSSDQPWPSSVTVAYGSQTGNAEGVAHVIAGQLRARGVAHVKCCEANDLEIGDLPAISHLVVVVSTAGQGEQPDNIKDFWKALENPALPADYLANTTFAVFGLGDSSYCFFCKSAIEIDERLGQLGAQRVLARGIGDDQDEDKYETGFSNWVPELYEALKLPEGKKETGVPNKHYKVVLGGKAHLDFDLTKQLAPTPILTEGCARHVRMLSNGRLTSLDYDRNIRHMVIDLEGSGLRYGVGDALAIYPRNAPDRVADFLGFMGLDPLECIDELQFIEAEGQTATKQKPPVPAHLPLAQLFTDVLDLFGRPSKRFYERLALYASDPAEKAALERVCERSAAGGGAFAKLLEETPTYADVIRMYPSTRAALSIEHLVDILPVMKPRYYSIASSPHHLPGKLELCIVVVDWKTKSGADKFGTCTGYLKDLLDFGADGKASYTIAATMKPSAGMCMPAKPEFPVICAGLGTGLAPMRAMVQERLHQIKAGKKIGETVLFFGCRFRAKDYLYGEEWEQALKDGNLTHLRVAFSRDQKEKIYIQNKIEQEPA
>Cp_TRINITY_DN25427_c0_g2::TRINITY_DN25427_c0_g2_i1::g.427::m.427 TRINITY_DN25427_c0_g2::TRINITY_DN25427_c0_g2_i1::g.427  ORF type:3prime_partial len:447 (+) TRINITY_DN25427_c0_g2_i1:50-1387(+)
MAGKHGMEEAASVPAAEHAAKKQAVSGPPPPSATGVDVEGWIKANMKPYDGDASFLAGPTARTTKVMDEFRRLFKLELEKGVLDVDAKTPSTLTSHGPGYIIKGEEVIVGLQTDAPLKRSIKPFGGVSMVKKACESYGYKLDPSVEKIFTEYRKTHNQGVFDIYDEEMRKARHDHIITGLPDAYGRGRIIGDYRRIALYGVDRLIQEKNNDKKKTPATTEENMRTREELSEQVRALQDLKKMALSHGPEYDIGRPAATAREAIQWTYFGYLGAIKQQDGAAMSFGRLDGFFDVYIERDLAEGRITEEQAQEMIDDLVIKLRMVRFLRTPEYNDLFAGDPTWVTMVLAGADGKGGHFVSRTSFRILQTLYNLGPAPEPNLTVLWAEGLPENYKRFCARVSIDTSSIQYENDDVMRPLYGTDYGIACCVSPMAMGKQMQFFGARCNLA
>Cp_TRINITY_DN3651_c0_g1::TRINITY_DN3651_c0_g1_i1::g.166867::m.166867 TRINITY_DN3651_c0_g1::TRINITY_DN3651_c0_g1_i1::g.166867  ORF type:5prime_partial len:450 (-) TRINITY_DN3651_c0_g1_i1:874-2223(-)
QPARGSTIRVAATATDAVPHWKLALEELDKPKDGGRKVLIAQVAPAVRVAIAESFGLAPGAVSPGKLAAGLRALGFDQVFDTLFAADLTIMEEGSELLHRLTEHLEAHPHSDEPLPMFTSCCPGWIAMLEKSYPDLIPYVSSCKSPQMMLAAMVKSYLAEKKGIAPKDMVMVSIMPCTRKQSEADRDWFCVDADPTLRQLDHVITTVELGNIFKERGINLAELPEGEWDNPMGVGSGAGVLFGTTGGVMEAALRTAYELFTGTPLPRLSLSEVRGMDGIKETNITMVPAPGSKFEELLKHRAAARAEAAAHGTPGPLAWDGGAGFTSEDGRGGITLRVAVANGLGNAKKLITKMQAGEAKYDFVEIMACPAGCVGGGGQPRSTDKAITQKRQAALYNLDEKSTLRRSHENPSIRELYDTYLGEPLGHKAHELLHTHYVAGGVEEKDEKK*
>Cp_TRINITY_DN26540_c0_g2::TRINITY_DN26540_c0_g2_i1::g.267191::m.267191 TRINITY_DN26540_c0_g2::TRINITY_DN26540_c0_g2_i1::g.267191  ORF type:complete len:582 (+) TRINITY_DN26540_c0_g2_i1:100-1845(+)
MLSAATARAAALAGAAAPRIASSIAPALARTLPSLAKRLMSAKTAGLKGRGSLFSPEREPQHVFKPANEIIRAAEIDAALERAKALAKDESVIRDILGRARDRALLKDHGMQPLGGGEYVQGLTVEEAAILLQVDSNNQELMQMLYDTAFAIKNLIYGNRIVLFAPLYIANYCVNSCTYCSFRAPNTSMPRAVLTDEELRQEVEALQRMGHRRLLLLTGESPRYTFDQFLNALKIASEVRTDPCGSIRRINVEIPSLSVSDFRRLKETNCVGTYTLFQESYHPETYARAHPAGPKSDYEWRVQTMDRAQTAHVDDVGVGVLYGLHDYKWETLAMLQHAQHLDQTYGAGPHTVSIPRMQPAEGAPDAMNIPHPVNDEDFKKLVAVIRCAVPYTGMILSTRENPEMRRQLLRLGVSQMSAGSKTDVGSYHQNQVKPGAKPVAEHEHFTAGPEPSAQPATCTPKEAKDAAGQFTLQDHRSLDDVVGDLLELGFVPSWCTACYRKGRTGEAFMKIAKKGDIQALCHPNALLTLEEYLIDYASERTREIGKKVLEVETGHIPSERAKHAFERKLKKIDEGQRDLYF*
>Gt_TRINITY_DN23406_c0_g1::TRINITY_DN23406_c0_g1_i6::g.204809::m.204809 TRINITY_DN23406_c0_g1::TRINITY_DN23406_c0_g1_i6::g.204809  ORF type:complete len:1961 (+) TRINITY_DN23406_c0_g1_i6:209-6091(+)
MAIGIGFLNGVRLASRGCTKLCSGAFSKSVSQTHFRSFSESVRGRTTPFGPYGGLINKEESADMNAERLENLGATGEGTPLYEKFIMERMARRAQFLAENKTLVNQSHDIEMIKNTTWSCIDGNQAAAHIAYALSEVSFIYPITPSSPMGEMVDEWAANGLKNIFGQTLSVTEMQSEAGAAGALHGSLKAGTFASTYTASQGLLLMIPNMYKIAGELLPCVMHVSARTLCAHALNIFGDHSDVMAARTTGWVMLASENPQMVMDQALVSHLATMDMRVPVLHFFDGFRTSHEVNKVRVINYDDIKKIFPWDSVKTHHDSALSPMNPHIQGTNQGPDVFFQAAEAANAIHNTIPAYMQKWSEKVAEITGRRYGFFSYEGHPEAENVIVIMGSGAVTVHETVKYLNSHQNQKVGVLKVRLFRPWITERFMAALPASVKRIAVLDRVKEVGAIGEPLFCDVCTTLHLSGMDHIKVYGGRYGLGGKDFTPGMVLSVFKNLEAAHPKTKFTVGIVDDVTNLSLNVEQEVDMLPSGTIQCLIYGLGSDGTVGANKSAIKTIAQNTDNFAQGYFEYDSKKSGGLTVSHLRFGPHPINAPYLVKNADYIGIHKESYLSRLDMLRNLKENGTVVINCTFGPEKVENMLPPRMKYQLASKKASLYLINAVKVGRETGMGKRINMVMQSVFFKLSRVLPFEKAIELLKKDVQKMYGKKGEEVVKRNWDAIDRSIENLVEVPVPAQWAQIPVTTNSAGKITSAAGMRSYSTYNGNGKTAPFSPALRSRSIWRTSNSPSRRYFSTRNLSSPAVSACGSKPGCGGCSKTPEEIRKSTLEFVDEIVEPVNKFEGNDLPVSAFVPGGRVPTGTSAFEKRGIALQVPRVDMDKCTQCNYCSLICPHAAVRPFLFTKAELEGAPEGIKKGSRKAIGGGVLDNYNFRVQVSPLDCTGCELCVRICPADALFFEETDKAVELEAENWNYAISVPNKGEEIDKTTVKGSQFQQPLLEFSGACEGCGETPYVKLLTQLLGDRLVVANATGCSSIWGASAPSFPYTVNARGEGPAWANSLFEDNAEFGLGMRRAFKQRRNQLMFLVDDALEDKTVPLSDKLRKLLSQYSVMRHEDKLDMLLPKGRSFYNQLREKIIPLLREEAKNHEKIQELWDQADIFGRHSHWIIGGDGWAYDIGFGGLDHVLASEEHINVLVLDTEMYSNTGGQASKSTPRGAQAKFAETGKLTAKKDLGQYAMTYKNVYVASICIHVNHQQAIRALLEAEAYPGPSLVICYSPCISQGFPMAEAIQQCRDAVESGYWPLYRYNPLLAEGGNNPFQLDSKKISGDLFTFLAHENRFASVMRRNPSYAQAHQDRLQRQIADRAKALNVLSLDELPASLKAAGPSESVTVLYGSETGNAEEQAKSLFADLKARGTSATLSSLDDFDFEELPNQSTVLVVISTCGQGEFPANSHKFWMKLSDPTLPMSFLEGIKFSVFGLGDSTYSLFCVAAERIDVRLAELGATRILNRGIGDDRDEDRYYTGWDNWTPQLWNALHVPQKPLERKIPKPAYKVTRTQGEATPSVSNDKLVPPGSNPLKLMENTLLTPEGYDRDIRHYVFKIKDTNVEYKVGDVLAIYPRNHVDQVEEFCKMYGLDPNEELNVVSTPEARNQIPEELNVRQLLQCVLDIFGKPNRRFYDTLSLFATDPAEKQKLELITSEDPDGKALYRELSHDMANHADVLKRFPSARPPLEQLMDMIPVIKPRSYSIASAPSMHPDEIELCIVAVDWEVPSTGEKRFGQCTSYLRTTKPGDTIMCSVKPSSIVLPEDNKAPLLMAGMGTGLAPWRALTQHRIALKQQGIDVGPCTIYYGARKGATEYLYREEFEKYEKMGVLRMVTAFSRDQPQKIYVQHRIREDYENVFRQLMKEQGSFYVCGSSRNVPEDIYNAMKEVMMLGGGMQEADAEAALASLKMDGRYTVEAWS*
>Gt_TRINITY_DN18509_c0_g1::TRINITY_DN18509_c0_g1_i1::g.292317::m.292317 TRINITY_DN18509_c0_g1::TRINITY_DN18509_c0_g1_i1::g.292317  ORF type:internal len:443 (+) TRINITY_DN18509_c0_g1_i1:1-1326(+)
CVASCPVAAITEKDAVQDVKDLLENNVDYARLKLGGLLEESPSSNDMPVQFGDENVGFEKKLDDLRKYVTVVQTAPAVRITISEAFGLPPGATTSGQLVTALKMLGFDYVFDTNFTADLTILEEGNELLERVAKGGPFPMFTSCCPAWINMVEKVYPDLIPNLSTCKSPQQMFGSLAKSYFADKVGRDPSEVKVVSIMPCVAKKDEASRENLRTEAGGPDVDYVLTTRELARLIKMHRPKISFQTLPDSAYDSPLGKSSGAALLFGTTGGVMEAALRTAYALTAPQSGASMPRINFEEVRGLSGIREATVDLHGTPLQVAIAHQGSNLRELVDKIRAGKAPAYHFVEMMACRGGCIGGAGNPKDAVDTELLQKRFASVYQGDAGLPLRTSHDNPEIMQVYSEFLGKPLGHKSHELLHTHYTPRHPRQAQLRLQQQQQQQQQQ
>Gt_TRINITY_DN57659_c0_g1::TRINITY_DN57659_c0_g1_i1::g.306971::m.306971 TRINITY_DN57659_c0_g1::TRINITY_DN57659_c0_g1_i1::g.306971  ORF type:5prime_partial len:169 (-) TRINITY_DN57659_c0_g1_i1:686-1192(-)
TTFSIAMAYQQNGGRLDPFVEGLEALETLQDGDRVLISEACNHNRITSACNDIGMVQIPNKLEAALGGKKLQIEHAFGREFPELESGGMDGLKLAIHCGGCMIDAQKMQQRMKDLHEAGVPVTNYGVFFSWAAWPDALRRALEPWGVEPPVGTPATPAAAPATAASGV*
>Gt_TRINITY_DN13833_c0_g2::TRINITY_DN13833_c0_g2_i1::g.66842::m.66842 TRINITY_DN13833_c0_g2::TRINITY_DN13833_c0_g2_i1::g.66842  ORF type:internal len:465 (-) TRINITY_DN13833_c0_g2_i1:3-1394(-)
HTFKPPHEIVREAEIHEALFVTAERAKDAALVQDLLHKAEDRALLKGGGVTPPQSGSEYVQGLSVLEAATLLNVDSGNKELMQMLFDTAFRIKERIYGNRIVLFAPLYVGSYCVNQCTYCSFRASNHAAPRGTLTNDQLVREVQALQRQGHRRLLLVAGESPRYTFEQTLDALKVVSEVHTEPCGRIRRINVEIPTLSVSDFRRLKATNCVGTYTLFHETYHRDTYKRVHPAGPKSDYNYRLLTMDRAQTAGVDDVGIGALFGLYDYRFEVLGMLQHAQHLDQTYGAGPHTVSIPRIQPALGAPDSLNGGPSPVSDADFMKLVAVLRCAVPYTGMILSTRESPQTRKMLLKLGISQMSAGSKTDVGSYSKDDAKGGLPESCQEESKDMLAQFTLQDHRTMDEIVHDLMEDGFVPSWCTACYRKGRTGEAFMKIAKKGEIHNFCHPNGLLTLQEYLIDYASERTR
>Gt_TRINITY_DN23046_c0_g1::TRINITY_DN23046_c0_g1_i4::g.290658::m.290658 TRINITY_DN23046_c0_g1::TRINITY_DN23046_c0_g1_i4::g.290658  ORF type:5prime_partial len:907 (-) TRINITY_DN23046_c0_g1_i4:141-2861(-)
LSSRHAVAMATRRTRMYIPASKRFIGNDSIKSIDEKIQRIQEAQAMFATYDQKTVDHIFRQVAHAANKQRVPLAKLAVEETRMGLMEDKVIKNGVVIETSVSAYADMKTCGIIDRDIERGLIKIATPVGPLACITPCTNPTSTVIIKSLFALKTRNAAIFLPHPRASACSHEAMRICRDAAVAAGAPPNVLDCVEHPTLELNRHVMDHPGIKLIVATGGPGMVKASYSTGKPALGVGAGNASVLVDDTADLKMAASSIVMGKTFDNGVICASEQSTVVIESIFEEFKSLLQKRGVHFVYGEERKKLGEFLIKNGAINPDIVGQSAQTIAQKCGIKIPSDAVVIATEASEVGPHEPFSFEKLSPVMTLYRAPTFNEAKDLCASIVHFGGEGHTAVIHSNDENRIAAFAAAMPAHHLFANVPSSIAAVGTAFNFNVPPSLTLGVGTTGGSSIATNMTPASLINVKLMASRQEHMEWMKTPPNLYFNRNCTREAIADLAKPYPGGKRDTRAMIVTDRAMVEFGYCAKVQQMLEQIGFKVAVFDDVTPDPTIECVRAGAAACEQFKPEVLIGLGGGSPMDAAKGIRVFYEHPEAKLEDVGARFIELRKRTCPFPDTGSKVHKLVCIPTTSGTGSEVTPFAVITEGTHKYPLFSYRMTPDVAIVDSAFCDNLPKSLVANAGVDALVHAIEAYVSVASNDFTKAHATRAVKLLFEYLPESYKNGTIRAREMVHHASTIAGIAFANSFLGITHSLSHQIGGAFHTPHGMTNAILMEHVIDYNAVDAPTRMGVYPQYTHPLAKQRYAELARAIGCTGNSDYELVQKFKQRISELKKSLDIPATFAEAGINKDAYYKVIDSLAEHAFDDQCTPANPRFPLVPELRHILECAYDGQPLSSRMYLEPKEADAAKTKA*
>Pp_TRINITY_DN25484_c0_g1::TRINITY_DN25484_c0_g1_i1::g.70261::m.70261 TRINITY_DN25484_c0_g1::TRINITY_DN25484_c0_g1_i1::g.70261  ORF type:3prime_partial len:765 (-) TRINITY_DN25484_c0_g1_i1:3-2294(-)
MEALLMPGPSMAAASRLMQTCSGNAAAIWMRSCVMMPMASAIVARRAHLRSLVTVERHAQKLMRTRLNDSSMHKGLYGTGIALQQRKRWPGLCSIPALATSGMAAQVQSQGELSKQVSLVINGRHVQVPEGASVLEAARKAGVYVPALCYHPNLEPAGTCRLCLVDILRNNPARGKKVASCATQAVDGMVVLTNTPELKDHVRSQIAMQRYRHPDRCQTCAANGQCEFQDLVGRYDVPPPPFVKPRVKPSLEQHGRAHANDTSSPALDLDFEMCVLCLRCVRACSELQGMNILGVVARGNEEVVAPVYSLDMKETECISCGACVASCPVAAITEKDAVQDVKDLLENNVDYARLKSGGLLEESPSSNDMPVQFGDENVGFEKKLDDLRKYVTVVQTAPAVRITISEAFGLPPGATTSGQLVTALKMLGFDYVFDTNFTADLTILEEGNELLERVAKGGPFPMFTSCCPAWINMVEKVYPDLIPHLSTCKSPQQMFGSLAKSYFADKIGRDPSEVKVVSIMPCVAKKDEASRENLRTEAGGPDVDYVLTTRELARLIKMHRPKISFQTLPDSAYDSPLGKSSGAALLFGTTGGVMEAALRTAYALTAPQSGASMPRINFEEVRGLSGIREATVDLHGTPLQVAIAHQGSNLRELVDKIRAGKAPAYHFVEMMACRGGCIGGAGNPKDAVDTELLQKRFASVYQGDAGLPLRTSHDNPEIMQVYSEFLGKPLGHKSHELLHTHYTPRHPRQAQLRLQQQQQQQQQQ
>Pp_TRINITY_DN25451_c0_g1::TRINITY_DN25451_c0_g1_i2::g.33184::m.33184 TRINITY_DN25451_c0_g1::TRINITY_DN25451_c0_g1_i2::g.33184  ORF type:5prime_partial len:475 (-) TRINITY_DN25451_c0_g1_i2:1576-3000(-)
CLKFMGIVMARYVLQRNMCVVESYTRTLRCKYATEAASSLGGISRVNLGIFGRMNAGKSTLMNFLTQQHTSIVDATAGTTADTKIALLEIHGIGPCKVFDTAGLDEQGALGAKKLQKTVQVAKECDVILIVHKSSQQSWRYAMDEAGVRSILDVAKARDKQVALVENVDQGQGDDGDGFTELDSWGGFPRIRVNPTRKACFEPFVAFLEKLHASKDRSTKVPLIPSKYLGKDKRLMMNIPMDAETPSGRLLRPQANVQEFALSYYTSTSAFRMDLAAGRSRDLHVKDQERRRFQEYVLREQPDLIVTDSQAMDLVYGWLSSPQFAHIALTTFSIVMLRKAGADLDYFIASLRAFDALKNGDRVLIAEACNHNRITQICEDIGTVQIPRHIRKKGCSAKQDITIDHAFGREFPDDELATYSLVVHCGGCMIDAQKMQARIEDCKDLGIPITNYGLLLSHVQSPGALSKVVAPFLS*
>Pp_TRINITY_DN25631_c0_g1::TRINITY_DN25631_c0_g1_i6::g.67692::m.67692 TRINITY_DN25631_c0_g1::TRINITY_DN25631_c0_g1_i6::g.67692  ORF type:5prime_partial len:517 (-) TRINITY_DN25631_c0_g1_i6:819-2369(-)
RRGKKWSVETENTHTFKPPHEIVREAEIHEALFVTAERAKDAALVQDLLHKAEDRALLKGGGVTPPQSGSEYVQGLSVLEAATLLNVDSGNKELMQMLFDTAFRIKERIYGNRIVLFAPLYVGSYCVNQCTYCSFRASNHAAPRGTLTNDQLVREVQALQRQGHRRLLLVAGESPRYTFEQTLDALKVVSEVHTEPCGRIRRINVEIPTLSVSDFRRLKATNCVGTYTLFHETYHRDTYKRVHPAGPKSDYNYRLLTMDRAQTAGVDDVGIGALFGLYDYRFEVLGMLQHAQHLDQTYGAGPHTVSIPRIQPALGAPDSLNGGPSPVSDADFMKLVAVLRCAVPYTGMILSTRESPQTRKMLLKLGISQMSAGSKTDVGSYSKDDAKGGLPESCQEESKDMLAQFTLQDHRTMDEIVHDLMEDGFVPSWCTACYRKGRTGEAFMKIAKKGEIHNFCHPNGLLTLQEYLIDYASERTRALGEQVVQQESETITSDRARKAYERKLKRINAGEHDLYF*
>Kf_TRINITY_DN21562_c0_g1::TRINITY_DN21562_c0_g1_i1::g.63888::m.63888 TRINITY_DN21562_c0_g1::TRINITY_DN21562_c0_g1_i1::g.63888  ORF type:5prime_partial len:638 (-) TRINITY_DN21562_c0_g1_i1:48-1961(-)
GDLFTFLAHENRFASVMRRNPSYAQAHQDRLQRQIADRAKALNVLSLDELPASLKAAGPSESVTVLYGSETGNAEEQAKSLFADLKARGTSATLSSLDDFDFEELPNQSTVLVVISTCGQGEFPANSHKFWMKLSDPTLPMSFLEGIKFSVFGLGDSTYSLFCVAAERIDVRLAELGATRILNRGIGDDRDEDRYYTGWDNWTPQLWNALHVPQKPLERKIPKPAYKVTRTQGEATPSVSNDKLVPPGSNPLKLMENTLLTPEGYDRDIRHYVFKIKDTNVEYKVGDVLAIYPRNHVDQVEEFCKMYGLDPNEELNVVSTPEARNQIPEELNVRQLLQCVLDIFGKPNRRFYDTLSLFATDPAEKQKLELITSEDPDGKALYRELSHDMANHADVLKRFPSARPPLEQLMDMIPVIKPRSYSIASAPSMHPDEIELCIVAVDWEVPSTGEKRFGQCTSYLRTTKPGDTIMCSVKPSSIVLPEDNKAPLLMAGMGTGLAPWRALTQHRIALKQQGIDVGPCTIYYGARKGATEYLYREEFEKYEKMGVLRMVTAFSRDQPQKIYVQHRIREDYENVFRQLMKEQGSFYVCGSSRNVPEDIYNAMKEVMMLGGGMQEADAEAALASLKMDGRYTVEAWS*
>Kf_TRINITY_DN63625_c0_g1::TRINITY_DN63625_c0_g1_i6::g.224372::m.224372 TRINITY_DN63625_c0_g1::TRINITY_DN63625_c0_g1_i6::g.224372  ORF type:internal len:424 (+) TRINITY_DN63625_c0_g1_i6:2-1270(+)
TSIRLRGVLRSAHLFLREISLPCAAQVGGNCRLAYASGAGEERSSAFAIRGKLIAQASFSCEGGRWSDDIEKYPSMGEKWSGAVKLVDLNDFIAPSQACVVSLKGQQQPADDQQGTEAQDGMVQIQARKADFPQVHKAATDEAIKVSLHDCLACSGCITSAETVMLQQQSTDEFVQRLSSSSQAVVVSLSPQSRASLATHYGITPLQAFRKLTGFLKSLGVKAVFDTSCSRDLSLLESCEEFVDRYKRAGGNSGTQRGDLPMLASACPGWVCYAEKTHGRFILPNISSTKSPQQVMGTLVKRFACKRLGLRPEDIYHVTIMPCYDKKLEAARDDFIFAVQREGSDTDTGQAGPRVTEVDSVLTSVEILDLLESRGVDFAALEEAPLDLMMSSVDEAGHVYGVRGGAGGYLETIFRHAARALFG
